# Distinct sequence patterns in the active postmortem transcriptome

**DOI:** 10.1101/293589

**Authors:** Peter A Noble, Alexander E. Pozhitkov

## Abstract

Our previous study found more than 500 transcripts significantly increased in abundance in the zebrafish and mouse several hours to days postmortem relative to live controls. The current literature suggests that most mRNAs are post-transcriptionally regulated in stressful conditions, we rationalized that the postmortem transcripts must contain sequence features (3 to 9 mers) that are unique from those in the rest of the transcriptome – specifically, binding sites for proteins and/or non-coding RNAs involved in regulation. Our new study identified 5117 and 2245 over-represented sequence features in the mouse and zebrafish, respectively. Some of these features were disproportionately distributed along the transcripts with high densities in the 3-UTR region of the zebrafish (0.3 mers/nt) and the ORFs of the mouse (0.6 mers/nt). Yet, the highest density (2.3 mers/nt) occurred in the ORFs of 11 mouse transcripts that lacked UTRs. Our results suggest that these transcripts might serve as ‘molecular sponges’ that sequester RNA binding proteins and/or microRNAs, increasing the stability and gene expression of other transcripts. In addition, some features were identified as binding sites for *Rbfox* and *Hud* proteins that are also involved in increasing transcript stability and gene expression. Hence, our results are consistent with the hypothesis that transcripts involved in responding to extreme stress have sequence features that make them different from the rest of the transcriptome, which presumably has implications for post-transcriptional regulation in disease, starvation, and cancer.

**ABBREVIATIONS:**

UTR: untranslated regions
ORFs: open reading frames
OP: overabundant transcript pool
CP: control transcript pool
FP: false positive
RBP: RNA binding proteins
ncRNA: non-coding RNA
miRNA: microRNA

## INTRODUCTION

Understanding regulatory circuits and how they influence transcriptional dynamics are important for comprehending the response of biological systems to stress such as starvation, disease, cancer and even death. Under stressful conditions, most (90%) mRNAs are regulated post-transcriptionally [1] -- presumably because it is more energetically favorable than regulation at the transcriptional level [2].

Two studies have recently shown that hundreds of transcripts increase in abundance in vertebrate organs/tissues in response to organismal death [3, 4]. These increases could be due to active transcription and/or post-transcriptional regulation of the nascent transcripts. Post-transcriptional regulation involves RNA binding proteins (RBPs) and non-coding RNAs (ncRNAs) [5, 6] that form complexes with RNA motifs and regions of secondary structure within the RNAs [7]. While the binding of RBPs to specific motifs in a transcript is well documented [8, 9, 10], the binding of ncRNA, in the form of microRNA [miRNA), circular RNA, or long ncRNA [lncRNA) to specific motifs within transcripts is less understood. Apparently, some mRNAs and ncRNAs act as “molecular sponges” that bind miRNAs preventing them from performing their functions. For example, miRNA-16 is sequestered by mRNAs encoded by the Tyrosinase-related Protein 1 (*Tyrp1*) gene [11]. As a consequence, miRNA-16 tumour-suppressor functions are lost and cell proliferation occurs [12]. Another “sponge” example is lncRNA encoded by the *Meg3* gene that binds miRNA-664 counteracting its inhibitory effect on production of alcohol dehydrogenase [6]. These are examples of two RNAs acting as molecular sponges, -- yet, not all of the functions of ncRNAs are known at this time [13] -- other roles have been suggested [14, 15, 16].

Our previous study revealed that some transcripts increase in abundance with postmortem time [4]. As a step forward towards better understanding of possible mechanisms responsible for these increases, our present study examined sequence features [i.e., short mers) that are over- or under-represented within these transcripts. We recognize that short mers are not the only sequence features responsible for these increases – we begin with short mers because they are easily identified. That said other more complex features are probably yet to be discovered.

We rationalized that some mers are over- or under-represented in these transcripts because they serve as binding sites for RBPs or ncRNAs involved in post-transcriptional regulation. To investigate this phenomenon, we examined the presence/absence/ frequencies of mers up to 9 nt in length and compared them to controls, which consisted of random draws of transcripts from the rest of the transcriptome (i.e., those not increasing in abundance in response to stress). The results show that several thousand mers are over-represented in the postmortem transcripts of the zebrafish and mouse.

Further examination of the frequencies of the mers show that some transcripts have more unique mers than others, and that the density of the unique mers varies by transcript and region (i.e., 5’UTR, ORF, 3’UTR).

## METHODS AND MATERIALS

A schematic overview of the experimental design for the study is provided in **Fig *1***.

**Fig 1.**
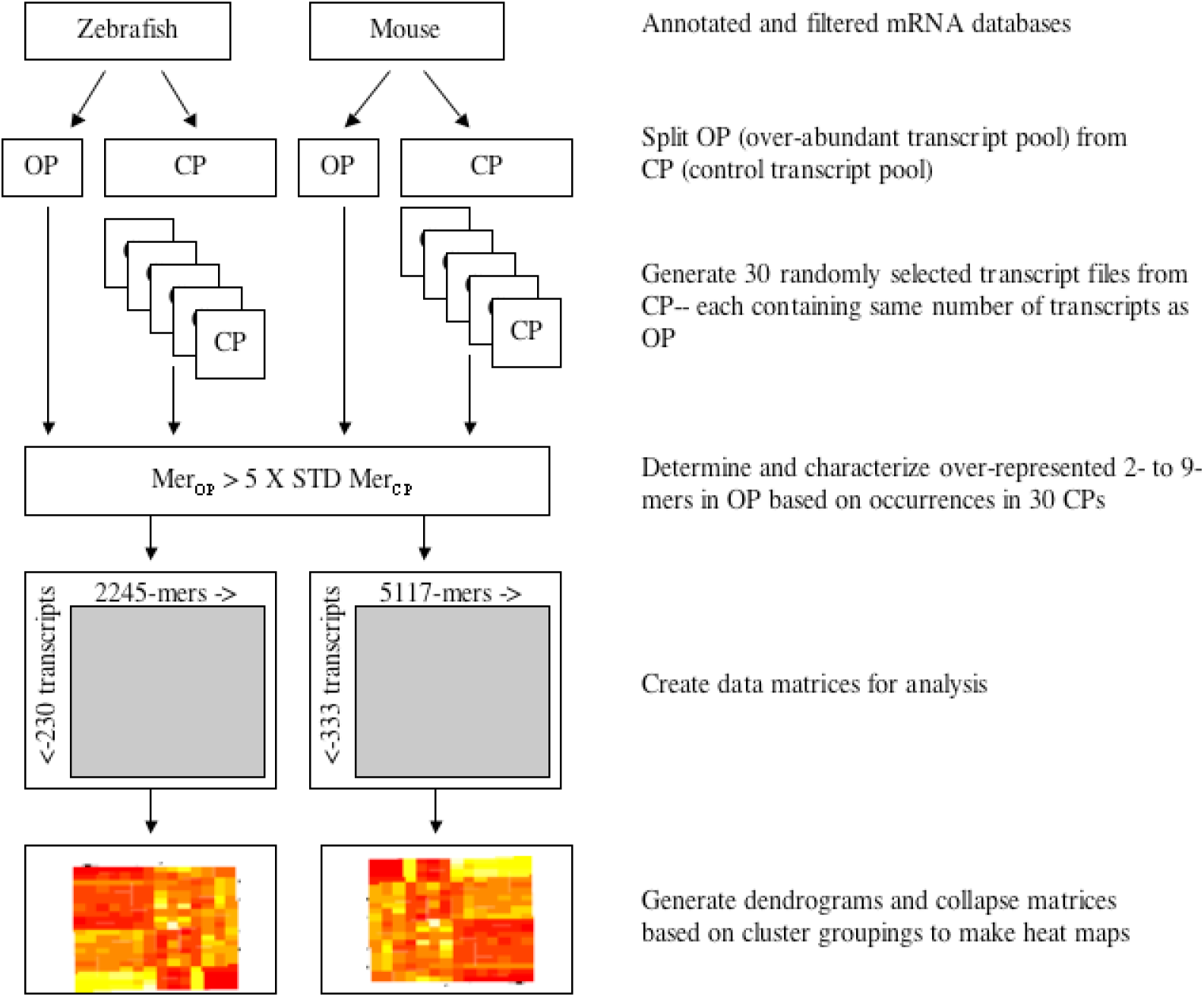
Schematic representation of the study experimental design.

### Dataset assembly

Messenger RNA transcripts of *Danio rerio* (GRC210.89) and *Mus musculus* (GRCm38.p5) with annotations were downloaded from NCBI. Transcript sequences containing ambiguous nucleotides (i.e., ‘N’s) and those less than 100 nt in length were removed. The final “clean” data sets were used for bioinformatic analyses.

### Extracting 2- to 9-mers from transcript sequences

An alignment-free sequence comparison method called ‘Chaos Genome Representation’ (CGR) [17, 18, 19] was used to extract mers from the transcript sequences because it was more practical (computationally efficient) than string-based search algorithms (see Proof in Online Resource 1). CGR is an iterative mapping technique that processes nucleotides in a sequence to find the *x-, y-* coordinates for their position in a continuous space. The *x-* and *y-* coordinates can then be used to recover sequence, which in this study were oligomers. Once the coordinates of a sequence are known, the presence/absence/frequency of a mer of any size in a transcript sequence can easily be determined, as demonstrated below.

### Reading a sequence into CGR space

The processing of a transcript sequence involves converting each nucleotide into *x-* and *y-* coordinates and assembling the coordinates into a CGR database. For example, the sequence ‘AAACC’ is represented by the *x*- and *y-* coordinates of +0.53125 and −0.53125, respectively. The coordinates are determined by reading the sequence into CGR space. The space is confined by the four possible nucleotides as vertices of a binary square with *x, y* position (−1, +1) being the vertex A, (+1, +1) being the vertex T, (−1, +1) being the vertex G and (−1, −1) being the vertex C. The position of a nucleotide in the fragment is calculated by moving a pointer to half the distance between the previous position and the current binary representation.

<u>An example.</u> Starting at point *x, y* (0, 0), the first nucleotide ‘A’ is plotted at half way to the vertex of A (−1, +1), which is coordinate (−0.5, +0.5). The next nucleotide is also ‘A’, therefore half way from the coordinate (−0.5, +0.5) to vertex of A (−1, +1) is (−0.75, +0.75). The next nucleotide is also ‘A’ so half way from the coordinate (−0.75, +0.75) to the vertex of A (−1, +1) is the coordinate (−0.875, +0.875). The next nucleotide ‘C’, so half-way from the coordinate (−0.875, +0.875) to the vertex of C (−1, −1) is the coordinate (+0.0625, −0.0625) and so on up to the last nucleotide of the sequence with the last coordinates of *x*=+0.53125 and *y*=-0.53125. A depiction of reading a sequence into CGR space is shown in Figure 1a of the Almeida et al. [19] study.

Once all the sequences have been read into CGR space and their coordinates stored in a database, it is possible to determine the presence/absence/frequency of mers by their coordinates and mer length (i.e., 1/resolution), which is outlined in the Mer analysis section below.

The software for the processing of nucleotide sequences into coordinates and recovering the sequences from the coordinates is available: http://peteranoble.com/software.html. Details on the mathematics of iterative mapping of nucleotide sequences have been previously published [19].

### Mer analysis

Mer analysis determines the presence/absence/frequency of a mer of length *z* (where *z* is 2 to 9) in a gene transcript.

<u>Finding a specific mer in a transcript.</u> Let us assume that a database of the *x-, y-* coordinates of the target sequence has been assembled and we want to determine the presence/absence of the mer ‘AAACAA’ in a target sequence. There are three steps.

First, we process the mer AAACAA into CGR space to find it *x-, y-* coordinates, which are −0.734375 and 0.734375, respectively.

Second, we determine the resolution of the search, which depends on mer length (i.e., resolution = 2^(mer length)^). A 6-mer requires a resolution of 64. The inverse of the resolution (1/resolution) is the CGR space around the coordinates that contain the specific mer. The CGR space around the coordinates is expressed by the following equation:

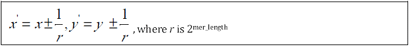

For the 6-mer AAACAA

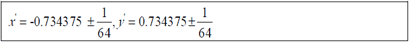

Third, the coordinates and CGR space of the mer is then used to search the CGR space of the target transcript sequence in the database. Any transcript that have coordinates within the box of x’ and y’ of the mer represents the sequence ‘AAACAA’. Furthermore, one can tally the number of hits within the box, which represents the frequency of the mer in a target sequence. We verified the presence of the mers in the identified target sequences by textual comparisons.

### Statistical and bioinformatics analyses

Analyses were conducted using SAS/JMP (version 7.0.2), R (version 3.4.0) and Microsoft Excel (versions 14.3.0 and 11.6). Hierarchical two-way cluster analysis was conducted on the binary matrices using Wards linkage method in SAS/JMP with default settings for cluster assignments. The resulting binary matrices were collapsed by their corresponding cluster assignments using a custom-designed program in C++. The resulting files were scaled to an average of zero and standard deviation of 1 in MS Excel and transferred to R to produce the heatmaps with no scaling. Network analysis was conducted using Gephi 0.9.2.

### Identification of 5’UTR, ORFs and 3’UTR and RNA motifs in transcripts

RegRNA 2.0 was used to identify functional RNA motifs and sites in the gene transcripts [20]. The server identifies splicing sites, splicing regulatory motifs, polyadenylation sites, transcriptional motifs, translational motifs, UTR motifs, mRNA degradation elements, RNA editing sites, riboswitches, RNA cis-regulatory elements, RNA-RNA interaction regions, and open reading frames using a integrated software package consisting of ~20 programs.

Nucleotide sequences of the transcripts were individually submitted to the server, default search parameters specified, and tab-delimited results downloaded to a computer. The results file contained global and local functions of the motifs and sites, their location in the transcript sequence, motif length and the sequence of the motif. Sequences of the unique mers in a transcript were matched to the sequence information of the motifs in the transcripts.

## RESULTS

Our previous study on postmortem gene expression dynamics [4] used a 60-mer oligonucleotide microarray to measure transcript levels. These perfectly matched probes were used in the present study to identify gene transcripts in the assembled datasets of the mouse and zebrafish (Online Resource 2). A certain portion of the transcripts has been shown to significantly increase in abundance after organismal death relative to live controls [4]. Henceforth, these transcripts are referred to as the over-abundant pool (OP), and transcripts not in this category are referred to as transcripts of the control pool (CP). Online Resources 3 to 6 contain probes and their corresponding transcripts. In total, the OP of the mouse and zebrafish consisted of 333 and 230 gene transcripts, respectively, and the CP consisted of 32,611 and 27,433 transcripts, respectively.

To determine if transcript length was a contributing factor when comparing different transcripts in the OP to those in the CP, we randomly selected two sets of transcripts from the CP (each set consisting of 333 gene transcripts for the mouse and 230 gene transcripts for the zebrafish) and compared the lengths of each set to those from the OP. No significant differences were found (two-tailed T-tests with unequal variance; alpha=0.05) in either the mouse or the zebrafish, which rules out transcript length as a factor affecting Mer analyses (Online Resource 7).

### Mer analyses

The occurrences of 2- to 9-mers in gene transcripts of the OP were compared to those of the controls (i.e., CP). In the zebrafish, the controls consisted of 2- to 9-mers found in 30 sets of 230 transcripts that were randomly drawn (with replacement) from the CP of the zebrafish (Online Resource 8). In the mouse, the controls consisted of 2- to 9-mers found in 30 sets of 333 gene transcripts that were randomly drawn (with replacement) from the CP of the mouse (Online Resource 9).

**Fig 2.**
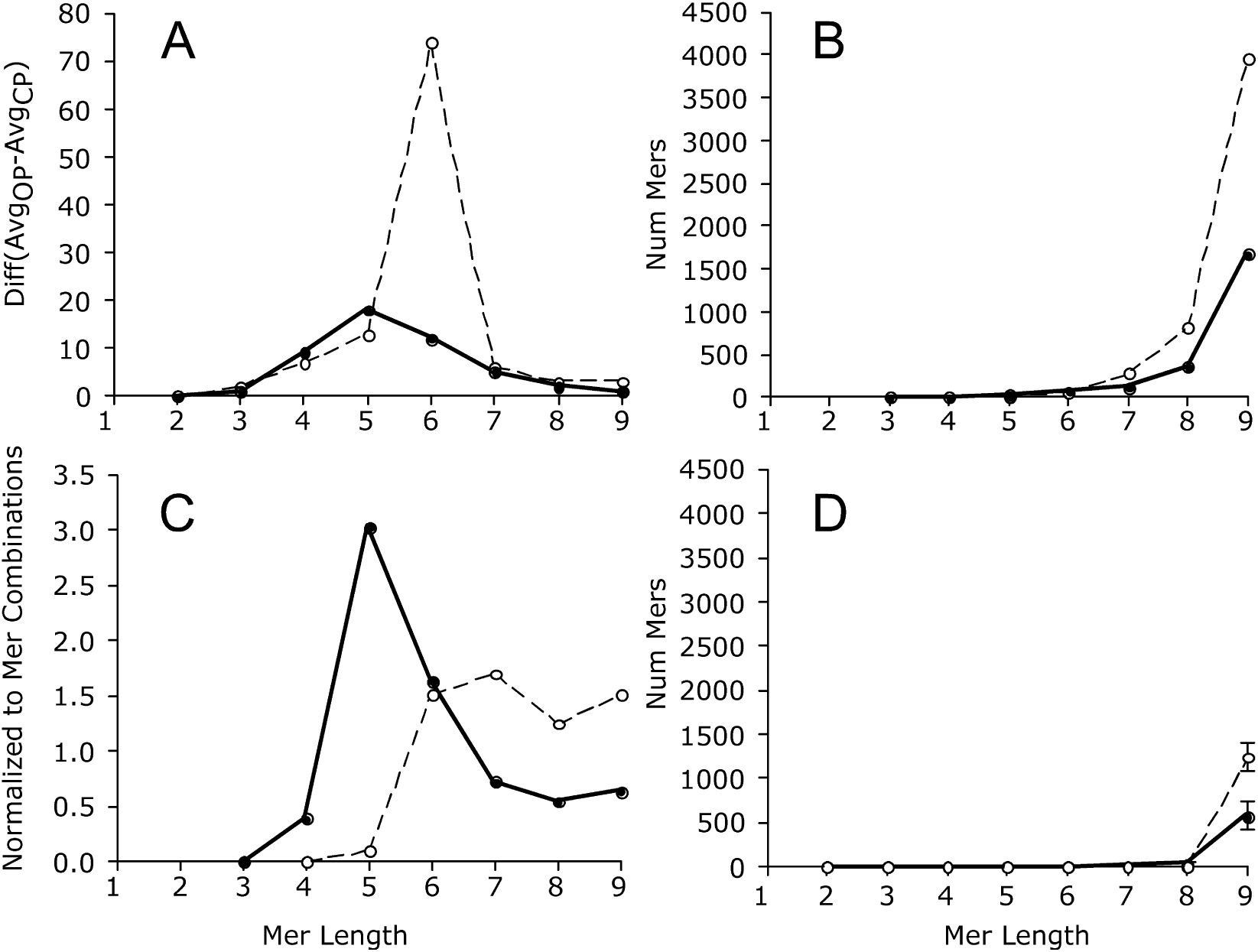
Mer counts as a function of mer length. Hatched line, mouse; solid line, zebrafish. Panel A, Difference in average mer counts by group (OP vs. CP); Panel B, individual mer counts that were 5 time stdev of average of the CP; Panel C, is the same results as panel B except normalized to the number of possible mer combinations and shown as a percentage; Panel D, number of mer counts that were 5 times stdev of the average CP due to random chance; average ± stdev of 3 random selections (without replacement).

To test the assumption that the 30 sets of random draws sufficiently represented the diversity of transcripts found in each organism, we classified an additional 3 sets of 333 and 230 transcripts from the CPs of the mouse and zebrafish, respectively (without replacement) (Online Resources 10 and 11). Only transcripts not previously drawn were used in this test.

The average count of individual mers from the random draws of the CPs were tabulated into a spreadsheet and compared to the counts of individual mers in the OPs of each organism. The arbitrary criterion used to identify ‘unique’ mers as either under- or over-represented was: a mer in the OP having less than or greater than 5 times the standard deviation of the average abundance of a corresponding mer in the CPs.

### Mer counts

Given that 2-mers have 16 possible nucleotide combinations (i.e., AA, AT, AC, … TT) and 3-mers have 64 combinations (i.e., AAA, AAT… TTT), all short mers (2 to 3 nt) were anticipated to be present in transcripts of the OP and CPs, and therefore, no differences between the pools should be observed. Differences between the pools however, should change with increasing mer length presumably due to real differences or random chance (i.e., false-positives; FP).

A maximum difference between the OP and CP pools was 6-mers (*n*=74 transcripts) for the mouse and 5-mers (*n*=18 transcripts) for the zebrafish (Table 1, **Fig *2*A**). When normalized to the number of possible mer combinations, the maximum difference was 7-mers for the mouse and 5-mers for the zebrafish (**Fig *2*C**). Hence, mers of 5 to 7 nt in length are optimal for distinguishing between the pools.

**Table 1.** Average ± standard deviation of 333 gene transcripts in the mouse and 230 transcripts in the zebrafish that contained unique mers by group (OP vs. CP). The absolute difference in unique mer counts by group and mer length is shown.

| Animal | Mer length | Num transcripts (OP) | Num transcripts (CP) | Absolute Difference |
| --- | --- | --- | --- | --- |
| Mouse | 2 | 333 ± 0.25 | 333 ± 0.4 | 0 ± 0.1 |
| | 3 | 330 ± 6.9 | 329 ± 7.9 | 2 ± 3.0 |
| | 4 | 304 ± 30.3 | 304 ± 30.4 | 7 ± 7.2 |
| | 5 | 227 ± 56.0 | 226 ± 55.9 | 13 ± 9.5 |
| | 6 | 119 ± 52.5 | 117 ± 50.7 | 74 ± 45.2 |
| | 7 | 43 ± 27.8 | 42 ± 25.5 | 6 ± 7.3 |
| | 8 | 13 ± 10.8 | 12 ± 9.1 | 3 ± 4.2 |
| | 9 | 3 ± 4.0 | 2 ± 1.9 | 3 ± 3.3 |
| Zebrafish | 2 | 230 ± 0.0 | 230 ± 0.1 | 0 ± 0.0 |
| | 3 | 230 ± 0.6 | 229 ± 1.8 | 1 ± 1.5 |
| | 4 | 220 ± 11.8 | 211 ± 15.5 | 9 ± 6.1 |
| | 5 | 166 ± 35.7 | 148 ± 33.5 | 18 ± 8.1 |
| | 6 | 81 ± 34.9 | 70 ± 29.4 | 12 ± 8.9 |
| | 7 | 27 ± 17.5 | 23 ± 14.2 | 5 ± 5.2 |
| | 8 | 8 ± 6.6 | 7 ± 5.0 | 2 ± 2.5 |
| | 9 | 2 ± 2.4 | 2 ± 1.5 | 1 ± 1.2 |

With increasing mer length, the number of ‘unique’ mers (i.e., over-/under-represented mers in the OP) increased (**Fig *2*B**).

To determine the number of FPs as a function of mer length and test the integrity of the experimental design, we randomly draw three additional sets of transcripts from the CP (without replacement) and retained only transcripts not used in the previous analyses.

For the mouse, each set consisted of 333 transcripts, and for the zebrafish, each set consisted of 230 transcripts. In this experiment, ‘over-/under-represented’ mers are FPs because the transcripts originated from the control transcript pool (i.e., the CP). To help explain the results of this experiment, let us consider the mer ATACCGG in the mouse. This mer would be considered ‘unique’ if its count were more or less than 5 times the standard deviation of the average from the CP, which is based on of 30 sets of 333 transcripts (Online Resource 11). The average and standard deviation in the CP was 8 ± 3.5, meaning one would expect to find it an average of 8 times in random draws of 333 mouse transcripts. Five times the standard deviation is 17.5, therefore the range of critical values for the mer count is: −9.5 and 25.5. In the OP, the mer occurred 31 times and is therefore considered ‘unique’ based on the stated criterion (i.e., the count is greater than 25.5).

To test the experimental design and check for FPs, mers were counted in three additional random draws from the CP. The mer ATACCGG, for example, occurred in 7 of the 333 transcripts in one set, 3 of the 333 transcripts in the second set, and 10 of the 333 transcripts in the third set (Online Resource 11). Since none of these counts are outside the criterion (the average ± standard deviation for this mer was 8.1 ± 3.49), there is no FPs for this mer. Of note, this procedure was repeated for all unique mers in the transcript pools of the mouse and zebrafish, respectively.

The results show that the number of FPs in the OP was close to zero for mer lengths of up to 8 bp (compare **Fig *2*D** to 2B). Therefore, while there is a possibility that some mers in the OP are FPs, the number was small (e.g., 8-mers: 1.0% are FPs in the mouse and 8.9% are FPs in the zebrafish).

When the length of mers was 9, however, the number of FPs significantly increased to an average (± std) abundance of 1240 ± 167.2 for the mouse (31.3% FP) and 571 ± 158.8 for the zebrafish (34.2% FP).

The results are consistent with the notion that unique mers can be identified in the OP by comparing them to random draws of mers from the CP. However, FPs increased with mer length. Taken together, over- and under-represented mers were identified in the OP and many are 5 to 7 nt in length.

### Survey of the unique mers

The survey of the OP identified 5,117 unique mers in the mouse and 2,245 mers in the zebrafish (Table 2). Normalized to the total number of combinations of 3- to 9-mers (*n*=349,504), this represents ~1.5% of the total mers in the mouse and ~0.6% in the zebrafish. Of note, 47 of the unique mers were common to both organisms (Table 3).

**Table 2.**
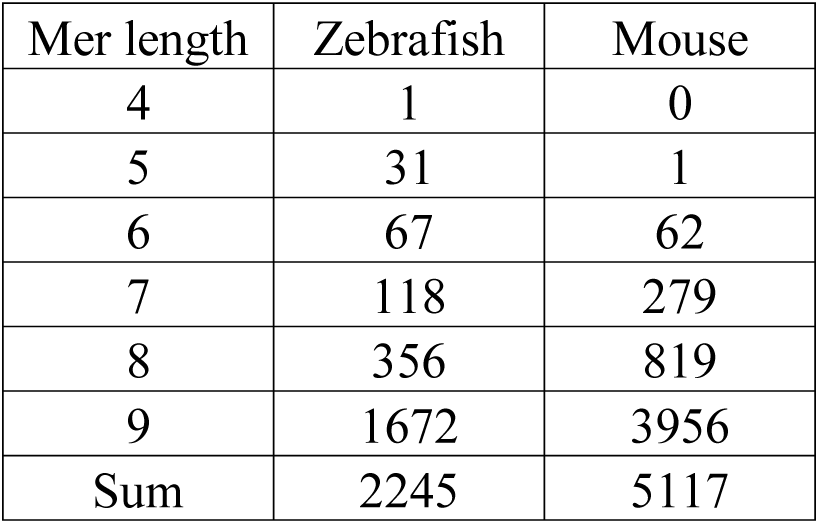
Number of unique mers in transcripts of the OP by mer length and organism.

**Table 3.**
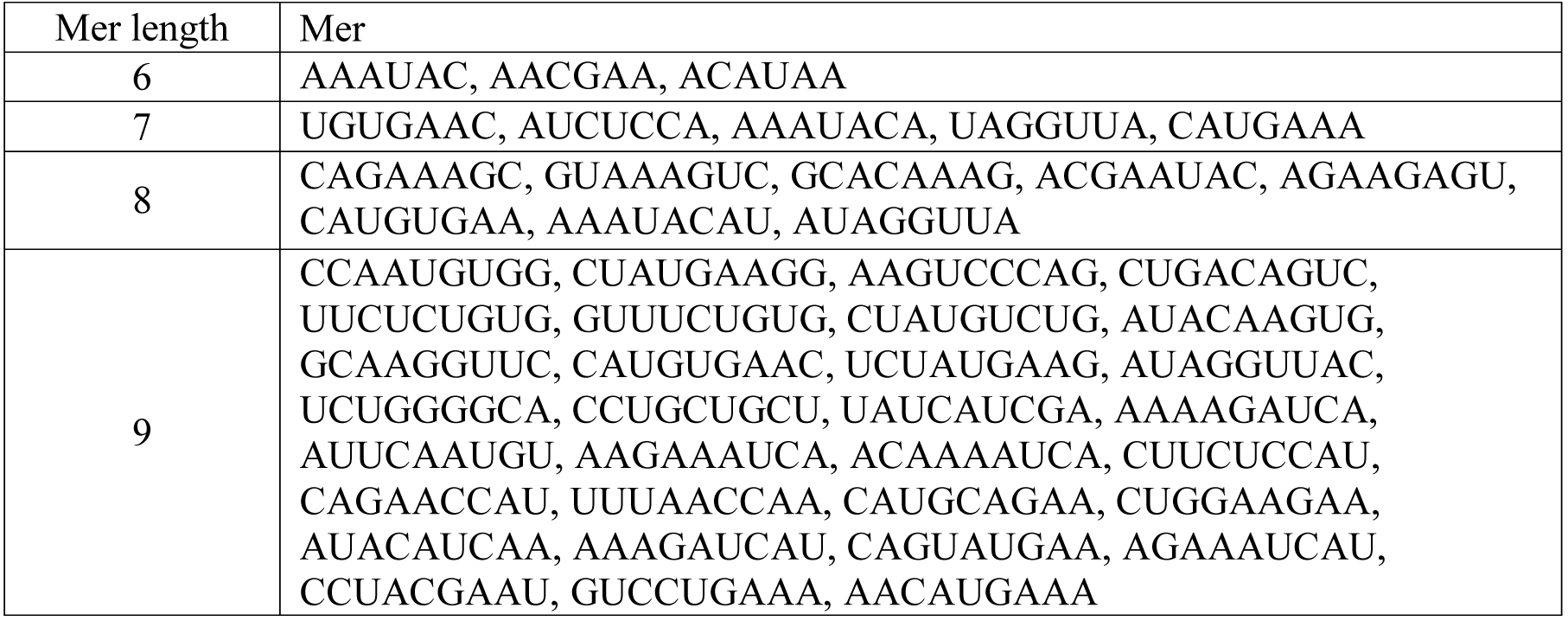
Unique mers common to transcripts of the OP for the zebrafish and mouse.

In fact, some of these mers are reverse complements to one another, which is of interest because they might form secondary structures and play roles in post-transcriptional regulation (Table 4). In the mouse, 218 of the 5,117 mers (4.3 %) reverse complemented one another. In the zebrafish, 31 of the 2,245 mers (1.4 %) were reverse complements.

**Table 4.**
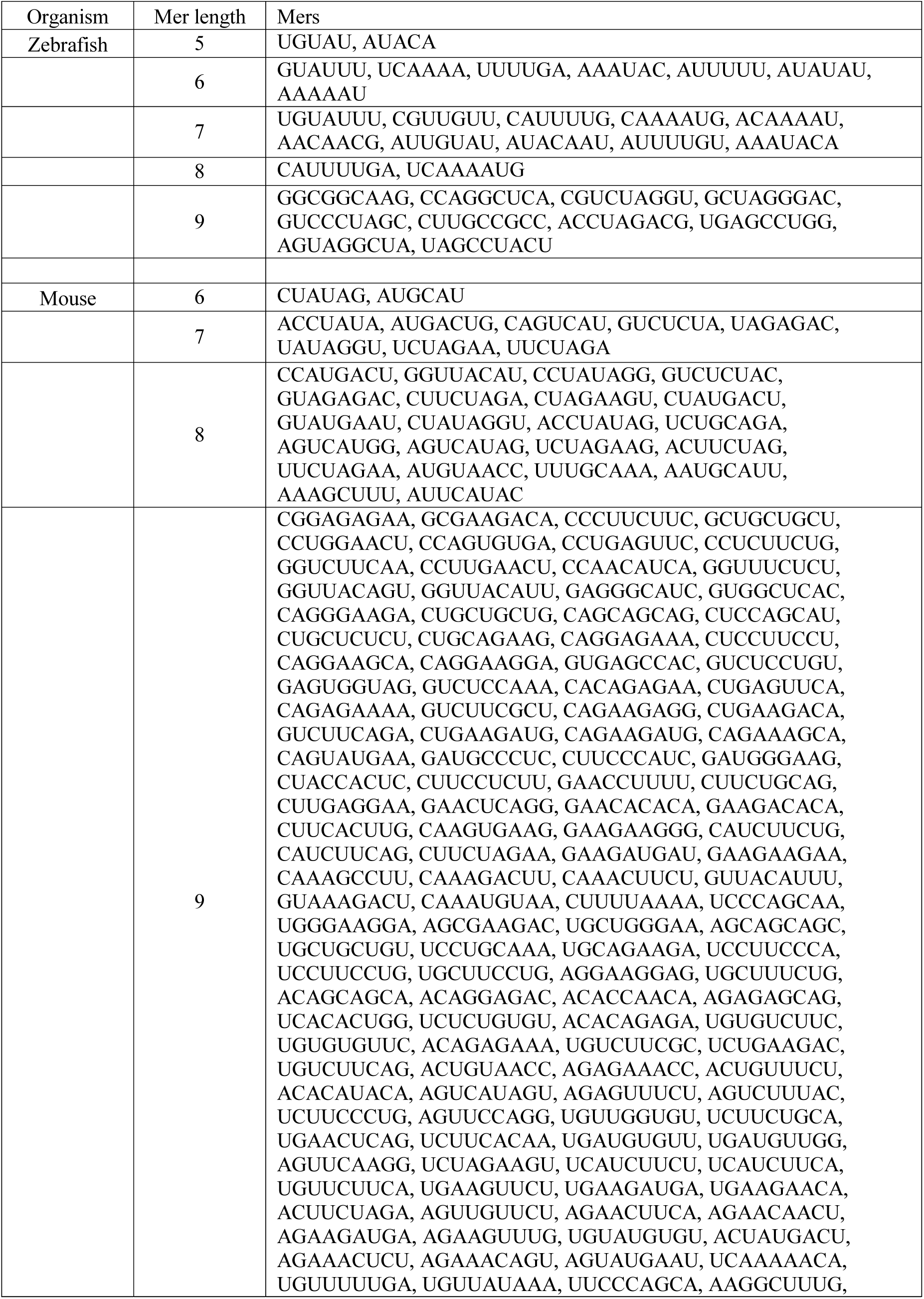
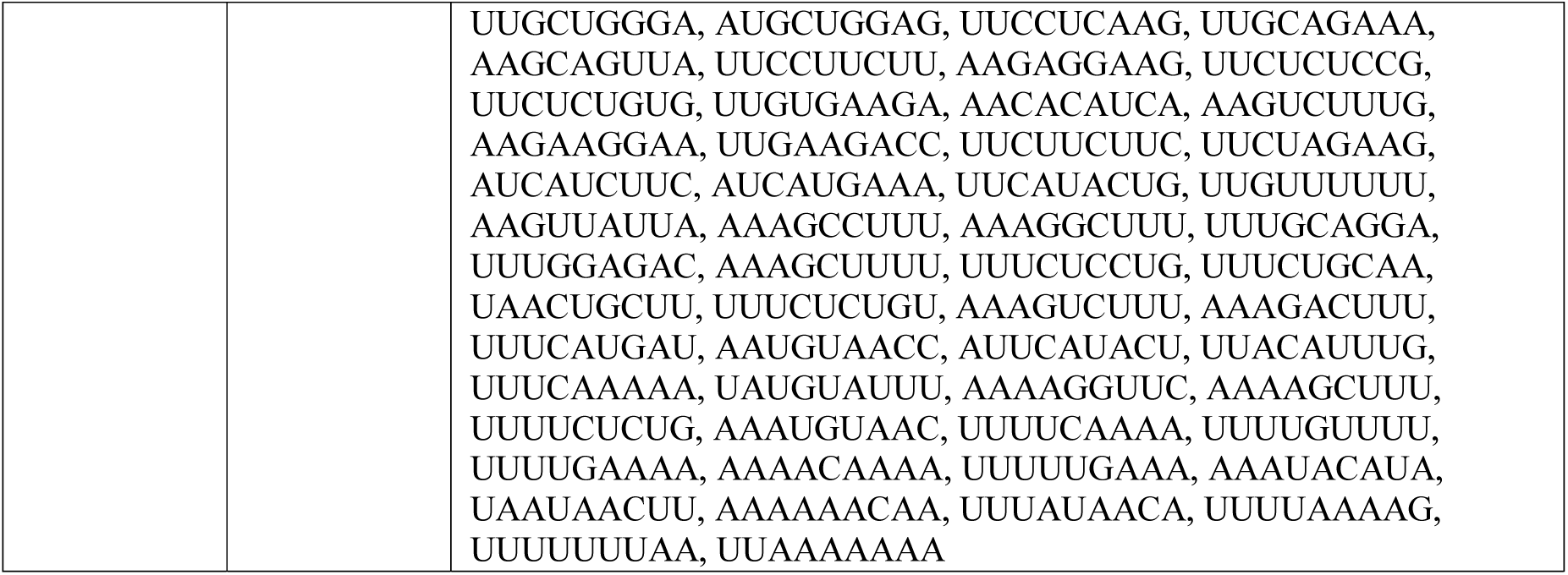
Unique mers that were reverse complements by length and organism.

### Number of unique mers per transcript

The distribution of the unique mers was investigated to determine if they were found in all transcripts of the OP, or just a few. In other words, *is the distribution of unique mers uniform across all transcripts?* To address this question, we compared their distributions in both transcript pools (i.e., OP and CP). Here we assumed that the corresponding unique mers in transcripts of the CP should approximate a skewed (Poisson) distribution because they are relatively rare occurrences. The controls in this experiment were the three sets of random draws (with replacement) from the CP. We also examined the multiple occurrences of unique mers in the OP since a unique mer might occur multiple times in the same transcript.

In the zebrafish, the frequencies of the unique mers per transcript varied between pools (**Fig *3***). These findings indicate that not all transcripts in the OP have the same number of unique mers – i.e., the number of unique mers in a transcript was not uniform. In the OP, the maximum bin was 150 while the maximum bin in the CP was 100. Some transcripts of the OP have more than twice the number of unique mers in the 200, 250, and 300+ bins than those of the CP. Therefore, some zebrafish transcripts in the OP have many more unique mers than others.

In terms of multiple occurrences of unique mers in the zebrafish, the distributions differed by pool also, with multiple unique mers occurring within the same transcript when compared to controls (**Fig *3*B**). For example, about 87 of the OP transcripts had more than 300 multiple unique mers compared to about 40 in the CP (**Fig *3*D**). Hence, not only are there many more unique mers in the OP but, in some cases, there are multiple occurrences of the same mer in the same transcript.

In the mouse, the frequency distribution of unique mers per transcript was also different between the pools (**Fig *4*A**). Specifically, there was almost double the number of unique mers in the 200 bin of the CP than the OP, about the same number of unique mers in the 400 bin, and twice (or more) the number in the 600, 1000, and 1200+ bins of the OP than the CP (**Fig *4*C**). This finding is consistent with those of the zebrafish – i.e., there are many more unique mers in the OP than the CP.

In terms of the multiple mer occurrences in the mouse, the results were different from the zebrafish; in general, there was little change between the histogram of the unique and multiple mers (compare **Fig *3*A** to 3b) – meaning that in contrast to the zebrafish, most of the unique mers did not occur multiple times in the same transcript sequence. Of note, this was not true for all cases as the 1200+ bin was somewhat bigger in the **Fig *4*B** than 4A. However, when compared to **Fig *3*B** to 3A, there is a substantial difference between unique and multiple mers in the zebrafish. The presumed reason for this disparity is that in the mouse, the unique mers tend to be longer in length than those in the zebrafish (Table 1, **Fig *2*C**) and the longer the length, the less frequent its occurrence.

**Fig 3.**
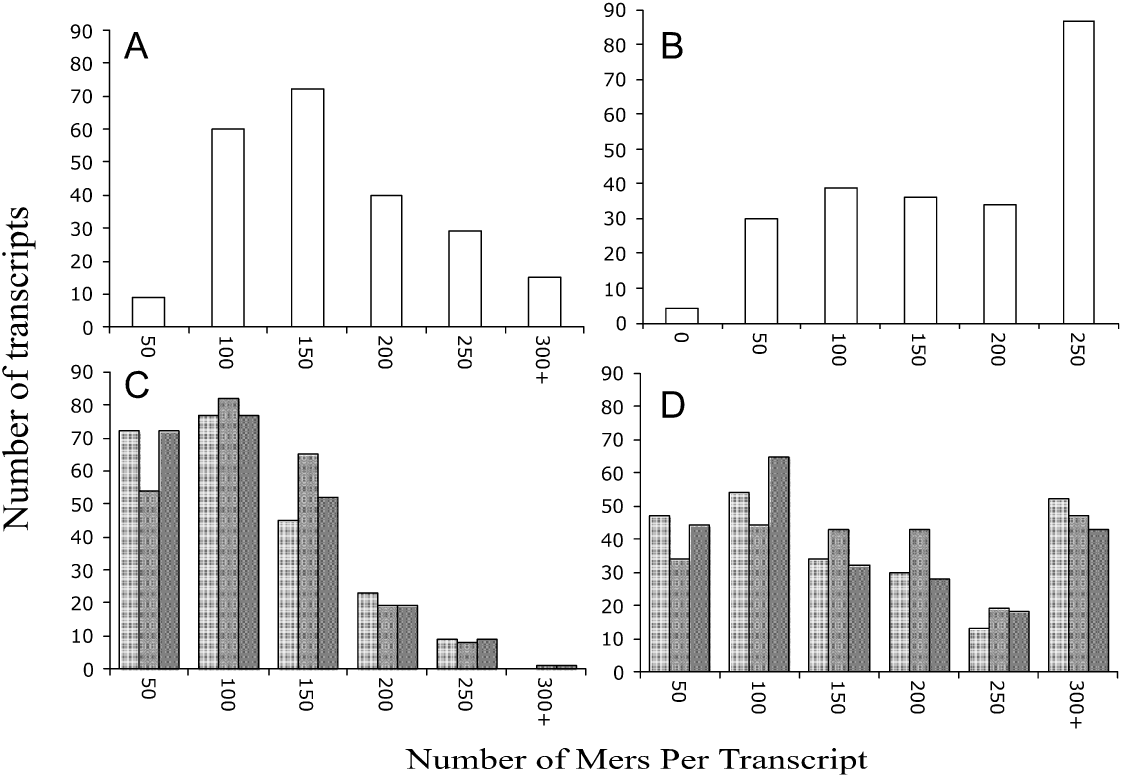
Distribution of unique mers per gene transcript in the zebrafish. A, unique mers in OP; B, multiple unique mers in OP; C, unique mers in CP (3 independent random selections; each as a different shade of grey); D, multiple unique mers in CP (3 independent random draws).

**Fig 4.**
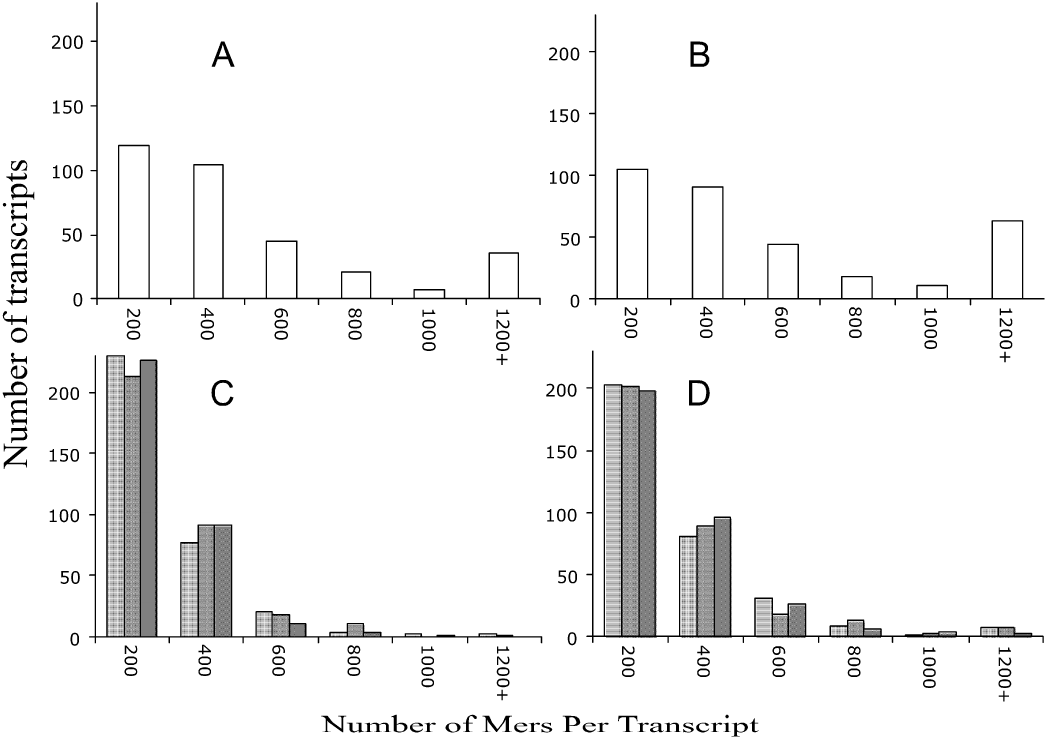
Distribution of unique mers per gene transcript in the mouse. A, unique mers in OP of the mouse; B, multiple unique mers in OP of the mouse; C, unique mers in CP of mouse (3 independent random selections; each displayed as a different shade of grey); D, multiple unique mers in CP of the mouse (3 independent random draws).

Taken together, the distribution of unique mers in the OP differs from those in the CP. Furthermore, there appears to be differences in multiple unique mers of these transcripts in the zebrafish but less so in the mouse.

### Groups of unique mers in the OP transcripts

Based on the previous analyses, we rationalized that some transcripts in the OP might share the same unique mers. To investigate the relationships among the OP transcripts and the unique mers (in binary presence/absence format), we constructed matrices and then performed two-way hierarchical clustering. The matrix for the zebrafish consisted of 230 rows of transcripts by 2245 columns of unique mers (Online Resource 12), and the matrix for the mouse consisted of 333 rows of transcripts by 5117 columns of unique mers (Online Resource 13).

The cluster analysis of the zebrafish identified 14 groups of transcripts and 20 groups of mers with high similarities, and the analysis of the mouse yielded 16 groups of transcripts and 20 groups of mers. The groups were collapsed by summation. For example, group A of the transcripts in the zebrafish consisted of 36 transcripts and Set 1 of the mers consisted of 25 unique mers. In total, 25 x 36 = 900 combinations, out of which 119 were actual occurrences of mers in the said transcripts (Online Resource 14), meaning there were 119 occurrences in the collapsed group. We summed groups A to N and mer sets 1 to 20 to form a collapsed matrix of 14 columns of transcript groups by 20 rows of mer sets. The same procedure was repeated for the mouse. The collapsed groups were normalized by row (see Materials and Methods section) to produce the data for the heat maps. Note, the heat maps were turned 90 degrees to show transcripts as columns and mer sets as rows.

The number of transcripts in a group and the number of mers in a set varied substantially for both organisms. Specifically, in the zebrafish, the number of transcripts in a group ranged from 1 to 59 (of the 230) (**Fig *5***), and in the mouse, the number of transcripts by group ranged from 1 to 124 (of the total of 333) (**Fig *6***). Hence, some transcripts are very similar to one another in terms of unique mers, while others are distinctly different – there was no uniformity (i.e., equal number of mers distributed to equal number of transcripts).

The number of unique mers in a set ranged from 2 to 876 (of a total of 2245) in the zebrafish (**Fig *5***) and from 40 to 1407 (of a total of 5117) in the mouse (**Fig *6***). Hence, some groups of mers are found in the same transcripts while others are found in different ones. Similar to the situation with the transcripts, the relationship among the mers was not straightforward– there appears to be a pattern.

There are unifying features visible in the heatmaps. For example, all transcript groups in the zebrafish contained relatively similar counts of mers within the mer sets 5 as well as 19 (**Fig *5***). Similarly, in the mouse, all transcript groups had similar counts of mers within the mer set 2 (**Fig *6***). Hence, despite similarities and differences of the collapsed data, there are common sets of mers found within all transcripts.

<u>Zebrafish heatmap</u>: In terms of differences, groups J and N are dissimilar from the other transcript groups (**Fig *5***) and each group consists of a single transcript. Group J represents the transcript *si_ch211-69b7.6*, whose function is currently not known, and

**Fig 5.**
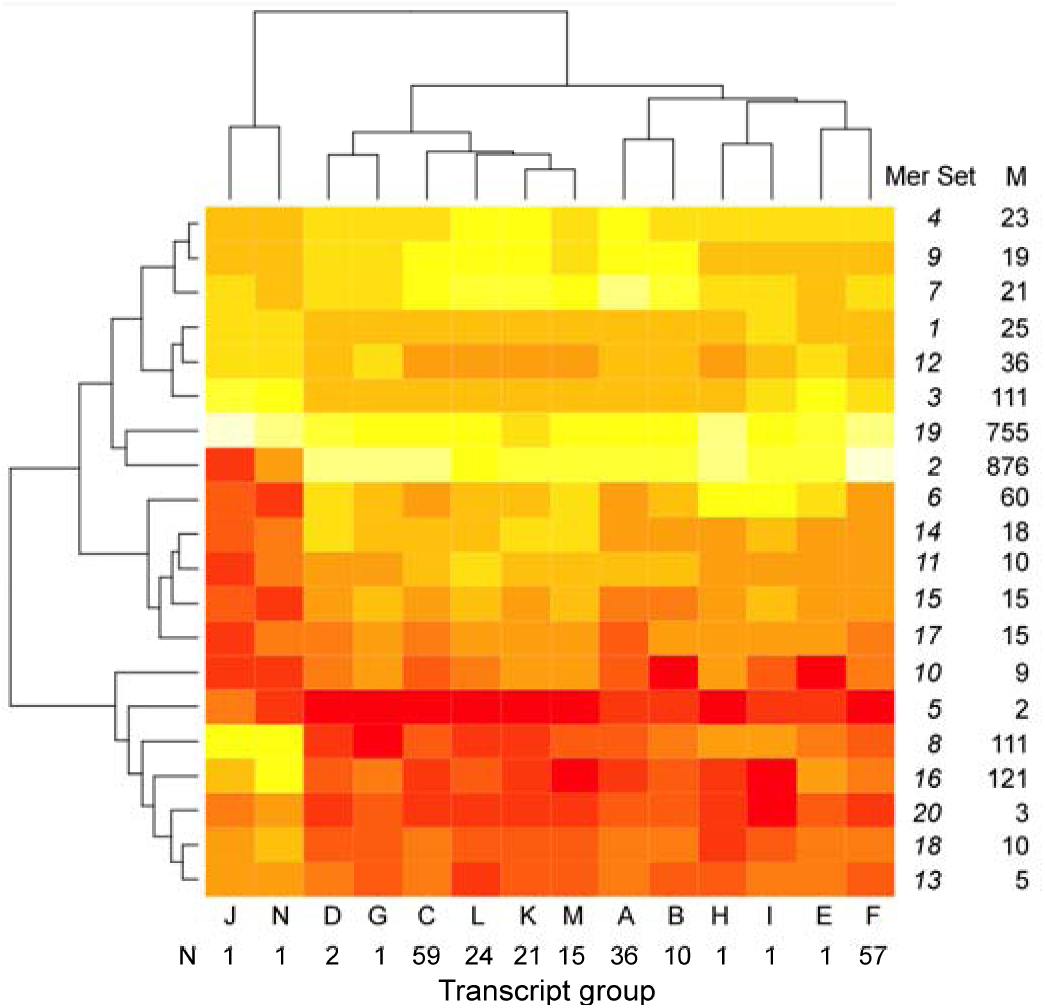
Heatmap of transcript groups and mer sets for the zebrafish. M, count of mers in group; N, count of transcripts in group. White, high count; yellow-orange, median count; red, low count.

**Fig 6.**
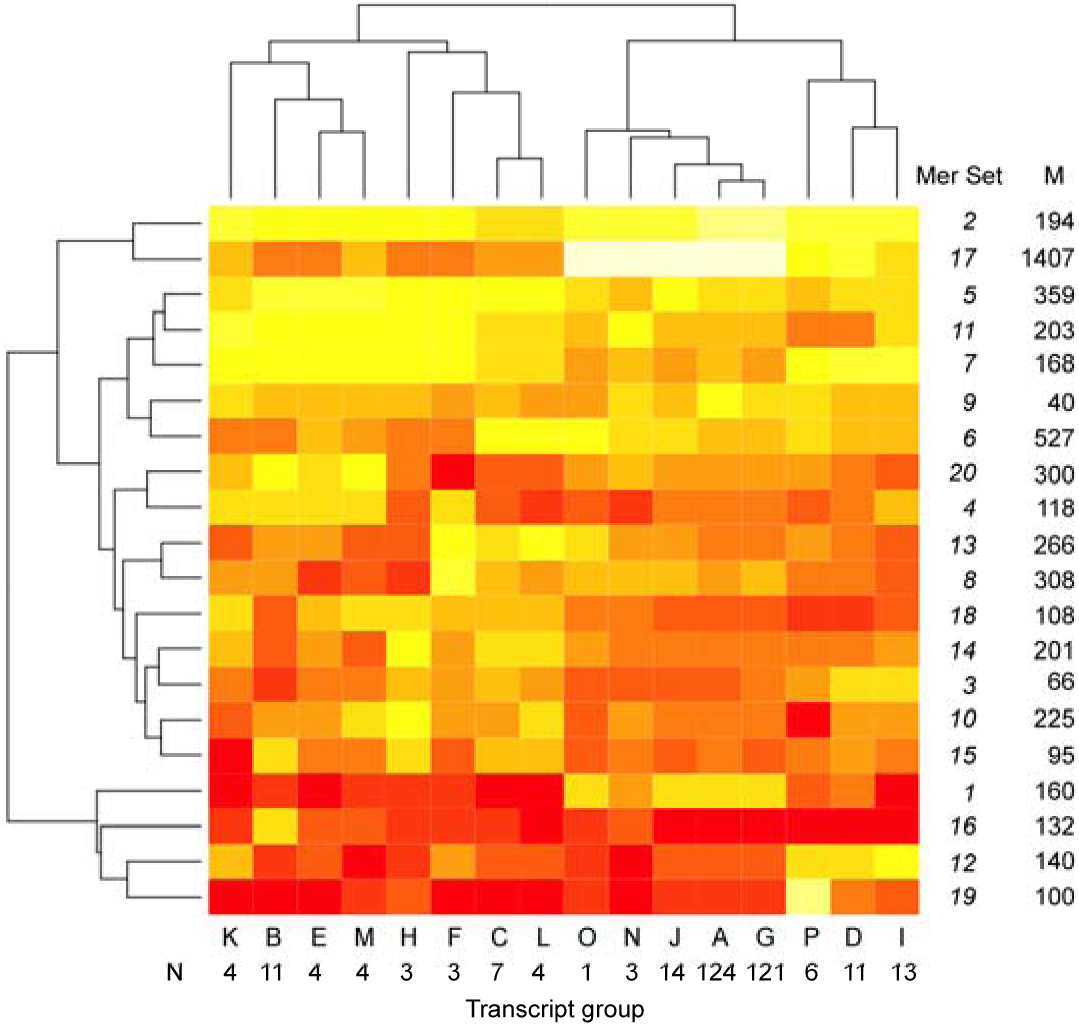
Heatmap of Transcript groups and mer sets for the mouse. M, count of mers in group; N, count of transcripts in group. White, high count; yellow-orange, median count; red, low count.

Group N represents the transcript *Psd2* (Pleckstrin and Sec7 domain containing 2), which is involved in regulating vesicle biogenesis in intracellular trafficking. The groups differed from the other groups in terms of the counts of mer sets 2, 3 and 8, which contain 876, 111, and 111 mers, respectively.

There appears to be significant differences between transcript group D, G, C, L, K and M, which consist of 122 transcripts (of the 230 possible) and group A, B, H, I, E and F, which consist of 106 transcripts (**Fig *5***). These groups are distinct due to subtle differences in mer set 10, which consists of 9 mers and mer set 6, which consists of 60 mers.

<u>Mouse heatmap</u>: The heatmap of the mouse shows similar variation in the number of transcripts by group and mer set (**Fig *6***). Transcript group K, B, E, M, H, F, C and L, which represent 40 transcripts (of a possible 333) is different from group O, N, J, A, G, P, D, and I (293 of possible 333 transcripts). The mer set responsible for this difference is mer set 17, which contains 1407 mers. Group O, N, J, A, G, P, D, and I have a higher relative counts than group K, B, E, M, H, F, C and L. Interestingly, the 40 transcripts in group K, B, E, M, H, F, C and L are annotated as either zinc finger proteins or predicted coding genes – and not one of the transcripts encode a protein with known function.

Taken together, there appears to be underlying patterns in the occurrence of unique mers in transcripts of the OP and these patterns are specific to certain groups of transcripts.

In the zebrafish, most (192) of the known functional gene transcripts are dispersed into many groups N, C, L, K, M, A, B, H, I and F, which represent 83% of the OP (**Fig *5***). In the mouse, most (245) of the known functional gene transcripts are found in groups A and G, which represent 74% of the OP (**Fig *6***).

### Density of multiple mers by transcript and organism

We examined the number of ‘unique’ mers by transcript length since longer transcripts might have more mers (Online Resource 15). Indeed, this was found true for the zebrafish -- there were more ‘unique’ mers with increasing transcript length (Pearson correlation coefficient, *r*=0.55, P<0.001). However, this relationship did not hold for the mouse (and we will show why below).

The averaged (± stdev) density of multiple mers for the zebrafish was 0.14 ± 0.18 mers/nt (*n*=230) and for the mouse was 0.40 ± 0.67 mers/nt (*n*=333). That is, there are 14 unique mers for every 100 nucleotides in the transcripts of the zebrafish and 40 mers for every 100 nucleotides in the transcripts of the mouse. Note the high standard deviations indicating a wide variation in values.

The highest and lowest densities of unique mers also differed between organisms. In the zebrafish, the highest density was ~1.0 mers/nt for *Pimr* gene transcripts, which corresponds to clusters B and H (**Fig *5***), and the lowest density was ~0.04 mers/nt for transcripts found in cluster A. We plotted the relationship between multiple mers and transcript length to find that the *Pimr* gene transcripts are distinctly different (red dots) from those in the rest of the transcripts in the OP (black dots) (**Fig *7*A**). The *Pimr* genes encode proto-oncogene serine/threonine-protein kinases involved in regulating the cell cycle. The remaining transcripts have a linear relationship between multiple mers and transcript length (*y*=0.1*x*; R^2^=0.91; with *x* is transcript length and *y* is multiple mers).

**Fig 7.**
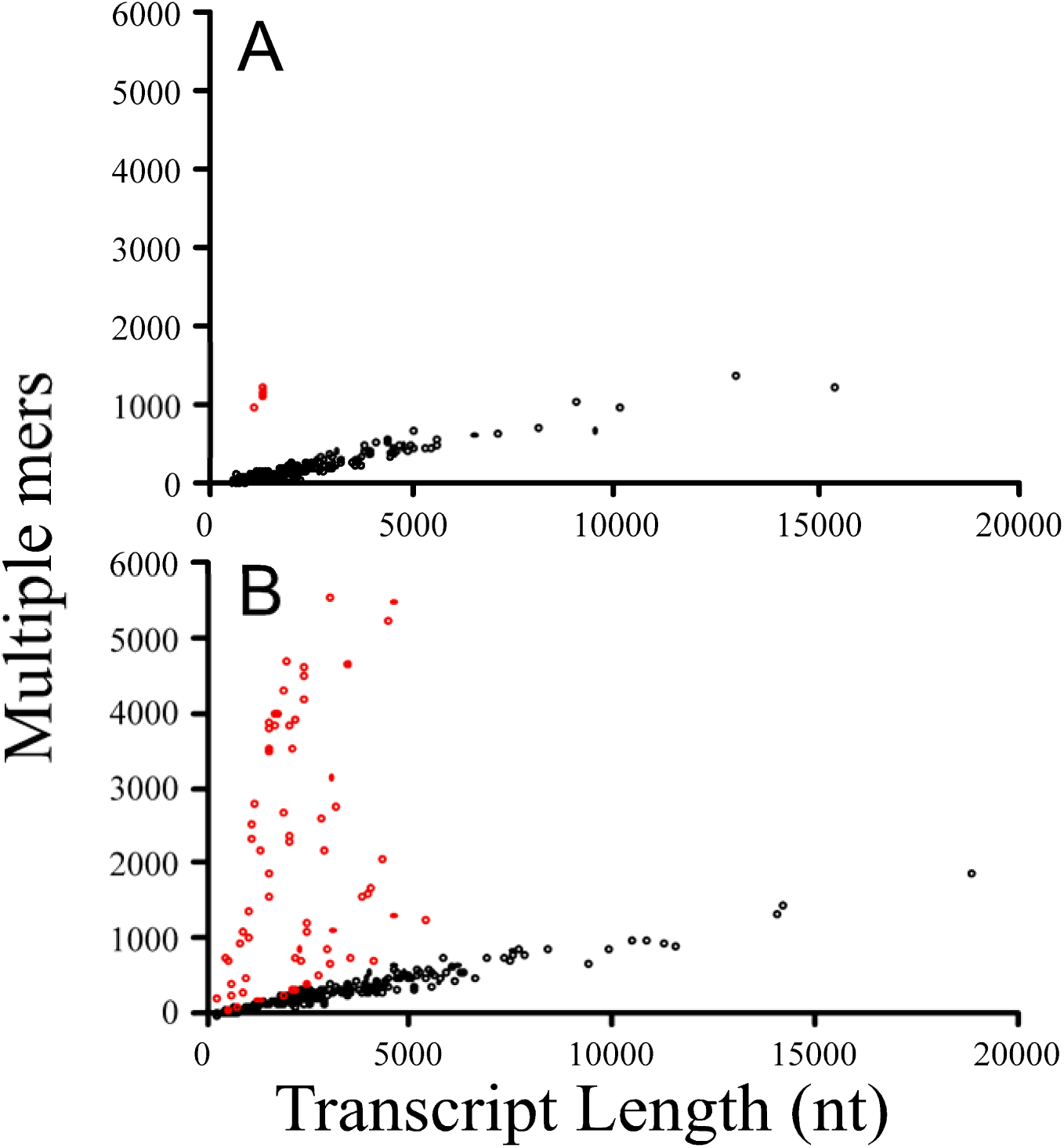
Number of multiple unique mers in transcripts versus transcript length. A, zebrafish; B, mouse; Red, deviant transcripts. Red dots in the zebrafish correspond to *Pimr* transcripts; Red dots in the mouse represent 47 transcripts (see text).

In the mouse, the highest density was ~2.6 mers/nt for annotated transcripts that do not have a canonical name (e.g., Gm14410, Gm14305. Gm14434, Gm2026, Gm11007, Gm2007, Gm4631) and were associated with Cluster B (**Fig *6***) and the lowest density was ~0.04 mers/nt in transcripts associated with cluster A. A plot of the multiple mers by transcript length for the mouse revealed significant differences for a subset of the transcripts (red dots) when compared to the rest (black dots) (**Fig *7*B**). The red dots represent 47 annotated gene transcripts, many that do not have a canonical name and includes those with the highest mer densities per transcript (mentioned above). The red dots also include 25 transcripts annotated as zinc finger proteins, 3 Rik transcripts, 1 unprocessed pseudogene, 1 Fam containing transcript, and 10 functional gene transcripts. The remaining transcripts have a linear relationship between multiple mers and transcript length (*y*=0.1*x*; R^2^=0.95; with *x* is transcript length and *y* is multiple mers). Hence the reason for the poor correlation between multiple mers and transcript length in the mouse data (noted above) was due to 47 transcripts that deviated from the other 286 transcripts in terms of their mer density.

We used RNAReg2 to determine if there are any unique molecular features in the 10 functional gene transcripts: *Bpifc, Ifitm7, Ms4a4c, Platr25, Rex2, Spag7, Styk1, Sva, Tmem239, Tnfrsf9*. We specifically looked at the relationship between the unique mers in the transcripts and the tab-delimited output files from RegRNA2 (Online Resource 15).

While all of the transcripts have ‘ncRNA hybridization regions’ that matched the unique mers, no patterns could be found in the AU-rich elements, K-boxes, UNR boxes, untranslated region motifs, long stem loop structures or transcriptional regulatory motifs among the 10 functional genes. Therefore, we concluded that the gene transcripts contain putative ncRNA hybridization regions – but we have no supporting evidence that these regions are actually used by the transcriptional regulation.

We rationalized that the transcripts with high mer densities might act as molecular sponges to RBPs and ncRNAs and thus alter their availability in the intracellular pools. If so, one would expect the profiles (i.e., transcript abundance by postmortem time) of transcripts with high densities and those transcripts affected by them to be highly correlated. Moreover, they should share similar unique mers that serve as putative binding sites. Principal component analysis was used to find patterns among transcripts with high mer densities using the correlations of their transcript abundance profiles to the rest of the profiles in the OP of the mouse brain. Network analysis was used to find shared mer binding sites.

The two axes of the ordination plot accounted for 96% of the variability (**Fig *8*A**). There appears to be three distinct areas in the ordination plot. One location is occupied by Gm14399, the other location is populated by a group of 8 gene transcript and the third location is occupied by Gm14409. The correlations among the transcript profiles differed by high density transcripts suggesting that certain groups might regulate different sets of transcripts.

**Fig 8.**
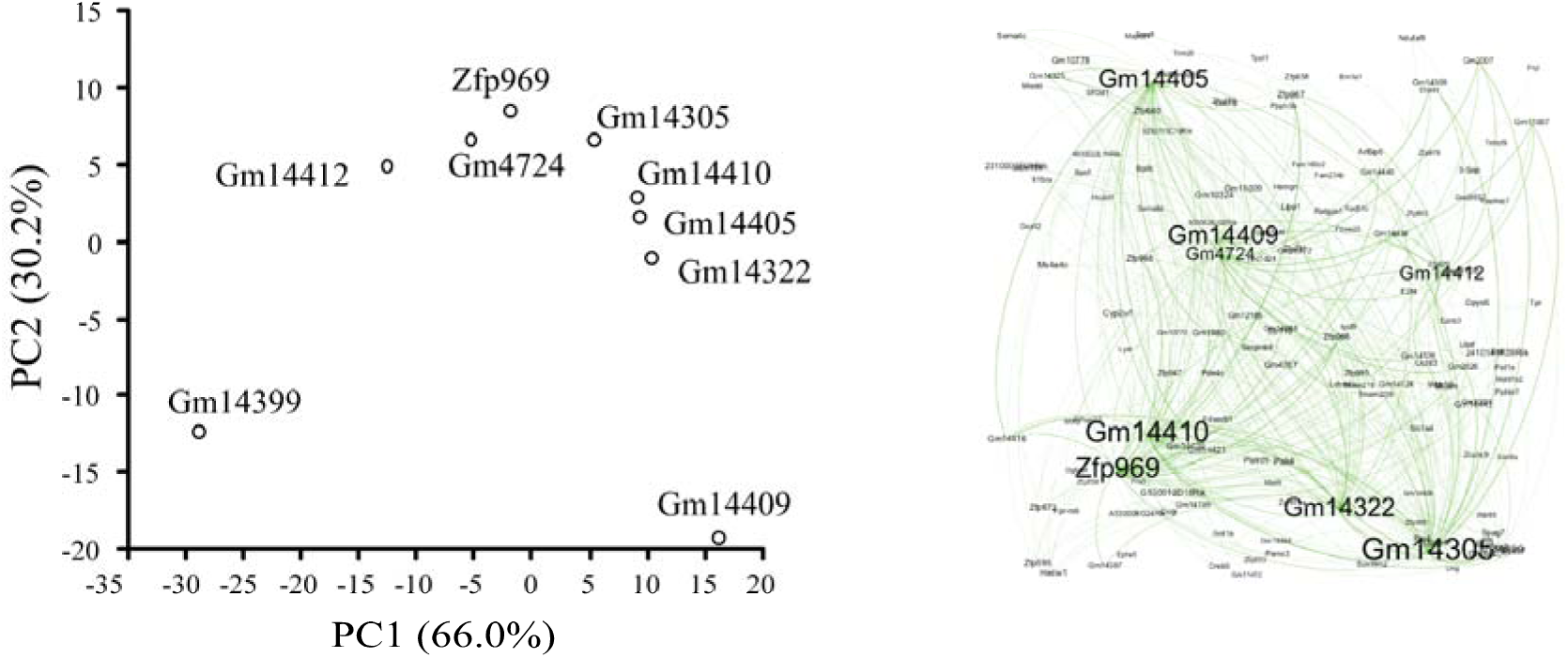
Ordination plot of transcripts with high mer densities (left) and network of transcripts with shared mers (right). The ordination was based on the correlations among mouse brain transcript profiles. The network was based on the number of shared mers in subset of the transcript profiles with high R^2^ (>0.95) to the transcripts with high mer densities. The network shows that the transcripts with high mer densities (i.e., molecular sponges) shared mers with many other transcripts.

To investigate the connections within the networks, we took a subset of the transcripts with high R^2^s (>0.95), and counted the number of shared mers. A network plot revealed that transcripts with high mer densities are connected to many different transcripts and that some shared similar mers. For example, Gm14305 shared mers with Gm11007, Gm2007, Gm14308 and *Hhmt1* as wells as many other transcripts (**Fig *8*B**). This finding suggests that the number of possible transcripts (and pathways) that are affected by molecular sponges appears to be quite vast.

Taken together, the results suggest that mer density is not the same in all OP transcripts and differs by organism and that transcripts with high density of mers have similar transcript profiles to the transcripts with lower density of mers some of which they share. The implications of this finding is that transcripts with high mer densities have the potential to act as molecular sponges to other transcripts and thus regulate them post-transcriptionally.

### Multiple mer density by region (5’UTR, ORFs, 3’UTR)

To investigate the density of unique mers by region, up to ten transcripts from each cluster (**Fig *5*** *and* **Fig *6***) were compared to determine if there are significant differences in mer density by region (Online Resource 16). Note that not all transcripts had 5’UTR and/or 3’UTR regions and some lacked ORFs (e.g., ncRNA).

In the zebrafish, for the transcripts having all three regions, the 3’UTR region had significantly more mers/nt than the other two regions (Table 5, Paired two-tailed T-tests, P<0.0001). Transcripts lacking 5’UTR, 3’UTR, or ORFs have low densities (i.e., ~0.1 mers/nt), indicating regional effects.

**Table 5.** Number of unique mers by nucleotide (transcript length), region and organism. Two-way paired t-test across rows: a,b, P<0.0001; c,d P<0.01.

| Organism | Regions | Number of transcripts | 5'UTR | ORF | 3'UTR | Non-coding |
| --- | --- | --- | --- | --- | --- | --- |
| Zebrafish | 5'UTR, ORF, 3'UTR | 70 | $0.2 \pm 0.26^a$ | $0.2 \pm 0.28^a$ | $0.3 \pm 0.35^b$ | - |
| | ORF, 3'UTR | 4 | - | $0.1 \pm 0.01$ | $0.1 \pm 0.03$ | - |
|  | 5'UTR, ORF | 1 | 0.1 | 0.1 |  | - |
| | ORF | 3 | - | $0.1 \pm 0.03$ | - | - |
|  | Non-coding | 1 | - | - | - | 0.1 |
| Mouse | 5'UTR, ORF, 3'UTR | 65 | $0.4 \pm 0.60^a$ | $0.6 \pm 0.70^b$ | $0.5 \pm 0.80$ | - |
| | ORF, 3'UTR | 2 | - | $0.1 \pm 0.00$ | $0.1 \pm 0.00$ | - |
| | 5'UTR, ORF | 16 | $1.4 \pm 0.80^c$ | $2.0 \pm 0.70^d$ | - | - |
| | ORF | 11 | - | $2.3 \pm 0.50$ | - | - |
| | Non-coding | 10 | - | - | - | $0.4 \pm 0.50$ |

In contrast, the highest unique mer densities in the mouse were found in the ORFs of transcripts – not the 3’UTR region as in the zebrafish (Table 5). In transcripts having all three regions, the ORFs had significantly higher densities than the 5’UTR (Paired two-tailed T-test, P<0.0001). In gene transcripts that have both 5’UTR and ORFs (no 3’UTR), or those having neither 5’UTR nor 3’UTR regions (i.e., ORF only) had twice the mer densities than transcripts having all three regions. Moreover, higher mer densities were found in the ORFs than the 5’UTR (Table 5, Paired two-tailed T-tests, P<0.01). One possible reason for these differences is that the 16 samples having no 3’UTR and the 11 samples lacking untranslated regions (i.e., they were all ORFs) consist of genes annotated as ‘predicted coding gene’ or ‘zinc finger protein gene’. Hence, gene function might play a role in these differences.

In summary, the results show distinct differences in mer densities by organism and region. In the zebrafish, the highest mer densities were found in the 3’UTR while the highest densities in the mouse were found in the ORFs.

### Known motifs

The following motifs are associated with increased mRNA stability or gene expression: the *Hud* binding site, YUNNYUY [21]; the *Rbfox* binding site, UGCAUG [10]; and UAUUUAU, GAGAAAA, AGAGAAA, UUUGCAC, AUGUGAA, UUGCACA, GGGAAGA [22]. We screened these motifs against the unique mers to identify transcripts in the OP that might have increased stability or gene expression due to these motifs.

Three hundred and fourteen of the 333 OP transcripts (94%) in the mouse and 189 of the 230 transcripts (82%) in the zebrafish contained one or more of the known binding motifs associated with increased mRNA stability or gene expression (Table 6). Most of the transcripts in the OP of the mouse and zebrafish had at least two different motifs (**Fig *9***). The number of previously reported motifs represents a small fraction of the total number of unique mers found in our study (180 of the 5117 unique mouse mers (3.5%) and 54 of the 2245 zebrafish mers (2.4%)). Hence, our study identified 4937 and 2191 putatively new motifs in transcripts of the OP of the mouse and zebrafish, respectively. It remains to be determined if these new motifs are functional or not.

**Table 6.** Number of transcripts by known protein binding site and organism. *Hud* binding site, YUNNYUY [21]; *Rbfox* binding site, UGCAUG [10]; and UAUUUAU, GAGAAAA, AGAGAAA, UUUGCAC, AUGUGAA, UUGCACA, GGGAAGA [22].

| Organism | *n* transcripts | Protein binding sites |  |  |  |
| --- | --- | --- | --- | --- | --- |
|  |  | *Hud* | *Rbfox* | Jacobsen et al. | All three |
| Mouse | 333 | 287 | 126 | 258 | 314 |
| Zebrafish | 230 | 185 | 0 | 106 | 189 |

**Fig 9.**
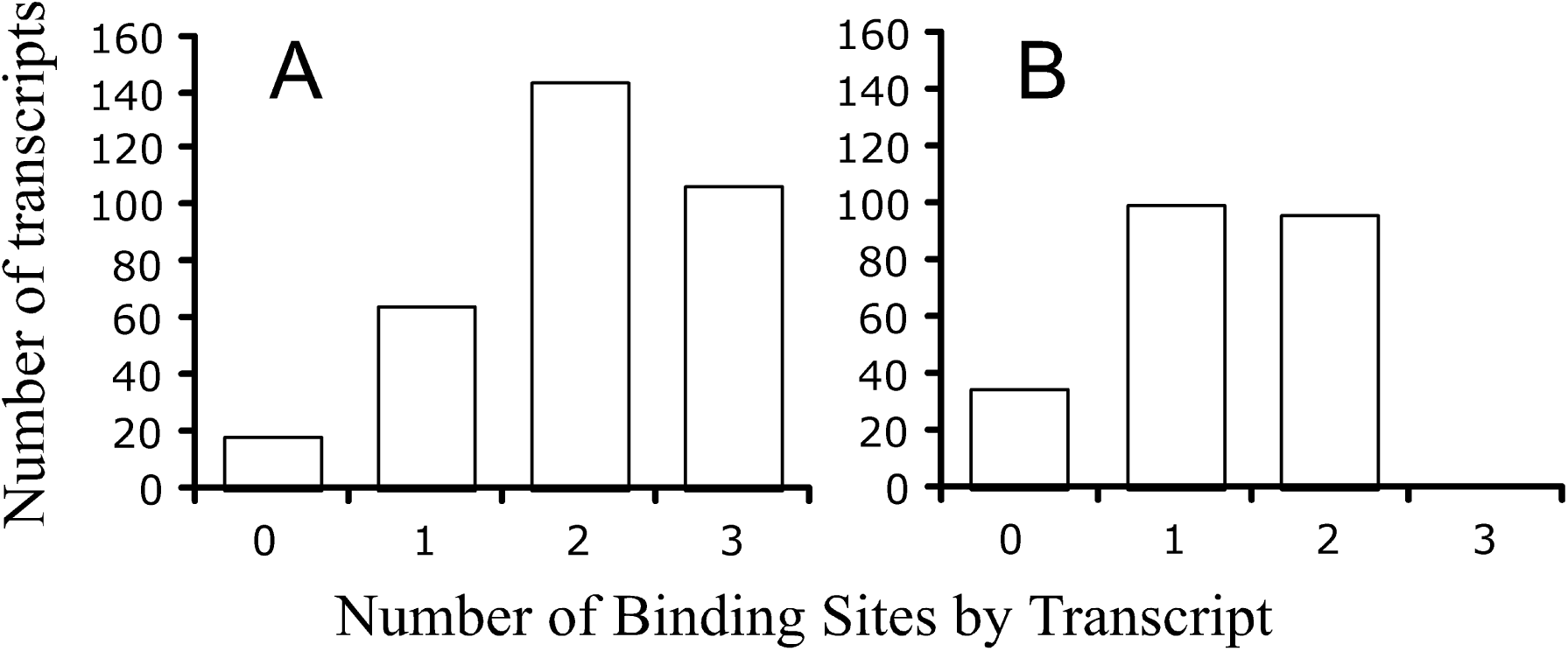
Number of known binding sites per transcripts for the mouse (A) and zebrafish (B). Total number of transcripts for the mouse, *n*=333 and for the zebrafish, *n*=230. The following binding sites were examined: *Hud* binding site, YUNNYUY [21]; *Rbfox* binding site, UGCAUG [10]; and UAUUUAU, GAGAAAA, AGAGAAA, UUUGCAC, AUGUGAA, UUGCACA, GGGAAGA [22]. Note: the zebrafish did not have *Rbfox* binding sites.

## DISCUSSION

The motivation for our study was driven by curiosity into possible mechanisms responsible for the increase in transcript abundances with postmortem time, which have now been reported to occur in the zebrafish, mouse, and humans [3, 4]. There is a need to understand regulatory features and how they influence transcriptional dynamics in order to comprehend the response of biological systems to stress. Yet, to our knowledge, no study has investigated possible reasons for increases in transcript abundance after organismal death. Such information is needed to provide baseline data for gene expression studies involving stressful conditions such as disease, starvation, and cancer.

### Unique mers identified in the OP

Our initial hypothesis was that among multiple reasons, there must be a signal, i.e., a nucleotide sequence that is responsible for postmortem activation of certain transcripts. Instead, we find sets of ‘unique’ mers in different groups of transcripts, with most sets consisting of ten to hundreds of different mers -- not just one or two.

The total number of unique mers in the OP was relatively small compared to all possible mers, ~1.5% of the total combinations of 3- to 9- mers in the mouse and ~0.6% in the zebrafish. These small percentages are presumably due to the arbitrary criterion used to identify unique mers. The reason the criterion was set to 5 times the standard deviation of the average count of the mer in the CP was to ensure that the identified mers were not due to random chance (i.e., false positives, FPs). Our results indicate that chance of a random mer having a count exceeding the criterion was relatively rare -- but FPs did occur and their occurrence increased with mer length (**Fig *2*D**).

The fact that several mers identified in our study have been previously reported to be involved with increased gene expression and/or mRNA stability (e.g., *Hud*, *Rbfox, ARE* binding sites; [10, 21, 23]) is consistent with the idea that our experimental design was effective at identifying ‘unique’ mers in the postmortem transcriptome of two different organisms.

### Unique mers by transcript, region, and organism

The number of unique mers in each transcript of the OP varied considerably. Some transcripts have a disproportionately high number of mers, while others have much lower numbers. Interestingly, in the mouse, several of the transcripts with high multiple mer densities have an ORF with no known function. Other transcripts have known functions, including: *Bpifc*, which is involved in innate immune response; *Fam160b2,* which is involved in phosphorylation of *Hsp70* [24]; *Ifitm7,* which is involved in regulation of cell proliferation and immune response [25]; *Ms4a4c,* which regulates receptor signaling and recycling [26]; *Spag7,* which is involved in antiviral and inflammatory response [27, 28, 29]; *Styk1,* which is associated with cancer progression and promotes the Warburg effect through signaling of the PI3K/AKT pathway [30, 31, 32]; and *Tnfrsf9,* which is involved in positive regulation of immune system functions and leukocyte activation [33]. In the zebrafish, a disproportionately high number of mers occurred in the *Pimr* gene transcripts, which are involved in cell cycling. These gene transcripts have common functions: cell survival, proliferation, cycling, stress compensation, and/or defense. It is enticing to speculate that the other transcripts (i.e., those with no known functions but with high mer densities) might also be involved in these functions.

The density of multiple unique mers was higher in the ORFs than the 3’UTR in the mouse -- but quite the opposite was true in the zebrafish (Table 5). That is, the zebrafish had a higher mer density in the 3’UTR than the other regions. In general, the 3′ UTR is involved in subcellular localization and mRNA stability, while the 5′ UTR play roles in translational control [34]. Motifs within the UTR regions are thought to control functions by interacting with RBPs [34]. Yet, the highest density of mers (2.3 ± 0.50 mers/nt) was in 11 transcripts that lacked UTRs (i.e., they were all ORFs). These findings are aligned with the notion that binding sites can exist all along the transcripts and not necessarily restricted to the UTRs [35]. It is possible that these 11 transcripts act as large “molecular sponges” in stressful conditions, providing an additional layer of complexity to post-transcriptional regulation (which we discuss below).

While the two organisms share 47 unique mers, there were significant differences in terms of their mer counts, the multiple mer densities by region, and the number of mers per transcript by organism. This finding suggests that post-transcriptional regulation varies significantly by organism – but this is not surprising since our original study [4] sampled mRNAs in whole organisms in the case of the zebrafish and the organ/tissues of the brains and livers in the case of the mouse. The samples are not comparable and we would not expect post-transcriptional regulation to be the same in different organisms or organ/tissues.

### Unique Mers and known binding sites

One set of unique mers with the sequence YUNNYUY apparently binds *Hud* proteins (Table 6). *Hud* proteins stabilize mRNA by binding to AU-rich instability elements (AREs) in the 3’UTR and they target transcripts involved in neuronal differentiation, protein phosphatase regulation, ubiquitin ligation, and the transport, processing and translation of mRNAs [21]. Interestingly, *Hud* proteins not only target their own mRNA but those of other RBPs, which suggests that it forms a network of post-transcriptional regulators [21]. In the mouse, data from our previous study [4] showed that *Hud* transcript abundance increased upon organismal death to reach maxima at 12 to 48 h postmortem (**Fig *10*A**). In the zebrafish, the *Hud* transcript abundance was about the same as the live controls for up to 4 h postmortem and then it declined and abruptly increased after 48 h (**Fig *10*B**). These findings are aligned with the notion that *Hud* genes are involved in stabilizing some of the mRNAs in our previous study.

**Fig 10.**
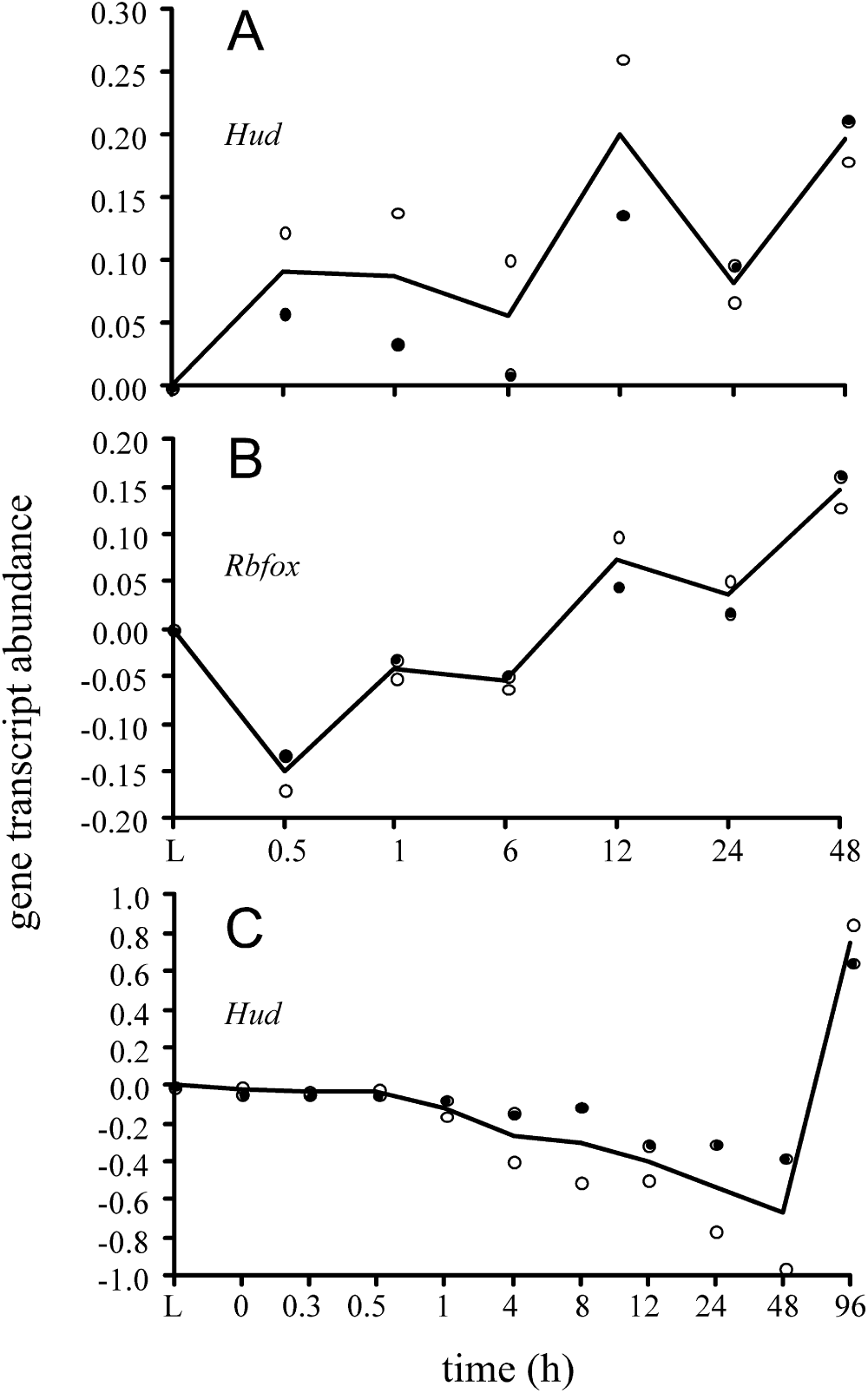
Gene transcript abundances measured by a calibrated microarray [41, 42] (log transformed) by postmortem time. Abundances were normalized to flash frozen live controls (L). Black line, average. (A) *Hud* transcript in mouse; black dots, averaged abundance measured by probe A_55_P1990309 (*n*=3 replicates for each dot except 48 h where *n*=2 replicates); white dots, average abundance measured by probe A_55_P1990314; (B) *Rbfox* transcript in mouse; black dots, average abundance measured by probe A_55_P195339` (*n*=3 replicates for each dot except last where n=2 replicates); white dots, average abundance of probe A_55_P1953400; (C) *Hud* transcript in zebrafish; black dots, average abundance of probe A_15_P119510 (*n*=2 replicates for each dot); white dots, average abundance of probe A_15_P120793. Data are from ref. [4].

Another unique mer with the sequence UGCAUG has previously been reported to serve as the binding site for *Rbfox* proteins that regulate splicing networks, mRNA stability and miRNA biogenesis [10]. Apparently, the binding to transcripts inhibits processing of the pri-microRNAs to pre-microRNAs, reduces expression of the mature miRNAs, and increases expression of targets normally downregulated by miRNAs [10]. A previous study has shown that the abundance of transcripts with UGCAUG motifs in the 3’UTR positively correlates with *Rbfox* expression, and that knockdown of *Rbfox* decreases transcript abundances [36]. These findings support the hypothesis that *Rbfox* enhances mRNA stability as well as gene expression. In our study, a little more than a third of the transcripts in the OP of the mouse have this binding site, but none were found in the OP of the zebrafish (Table 6). In the mouse, data from our previous study [4] showed that *Rbfox* transcript abundance increased after 30 min postmortem to reach a maximum at 48 h (**Fig *10*C**). These findings suggest that *Rbfox* proteins were interacting with some of the mouse mRNAs in our previous study.

The following 7 unique mers found in the OP have recently been reported as putative binding sites: UAUUUAU, GAGAAAA, AGAGAAA, UUUGCAC, AUGUGAA, UUGCACA, GGGAAGA [34]. These sites have been correlated with increased gene expression in HeLa cells transfected with miRNAs. The UAUUUAU binding site is reported to be an ARE that signals rapid degradation or increased stability of mRNAs in response to stress [36]. The Jacobsen et al. [34] study showed that ARE binding sites and miRNA mediated regulation are interlinked, which is aligned with a similar study in *Drosophila* cells [37]. While the significance and mechanistic insights of the 6 other putative binding sites were not discussed in the Jacobsen et al. study [34], at least one of the seven binding sites was found in 258 of the 333 transcripts of the mouse and 106 of the 230 of the zebrafish, indicating that miRNAs might be involved in “regulating” the postmortem transcriptome (Table 6).

### Post-transcriptional regulation of the postmortem transcriptome

Several possible scenarios could be working in spatially and temporally combination to increase transcript stability and/or increase transcript abundance in the postmortem transcriptome. These scenarios are based, in part, on the “Competing endogenous RNA hypothesis”, which is provided at the end of the Discussion. However, without experimental evidence, we caution that these scenarios are speculative at best.

One scenario is transcript stability is increased in the OP because they have more unique mers than the CP and RBPs bind to regulatory sites of transcripts of the OP blocking the binding of miRNAs, which are linked to degradation pathways. As a consequence, transcript stability is increased because the transcripts accumulate in the cells over time.

A second scenario is postmortem genes are upregulated due to miRNA inhibition. Take, for example, transcripts regulated by p53 tumor suppressor that increase in abundance in response to miR-21 inhibition [38].

A third scenario is that some of the transcripts containing high multiple densities of mers act as molecular sponges that bind miRNAs and/or RBPs and therefore affect post-transcriptional regulation *in trans*. An example of this in our study was the 11 gene transcripts in the mouse with unknown functions and the *Pimr* transcripts in the zebrafish that had high densities in terms of mers per nucleotide (~2.4 mer/nt and ~1.0 mer/nt, respectively). Such high densities indicate that they contained many unique binding sites to sponge RBPs and/or ncRNAs. According to the data from our previous paper [4], all the transcripts with high mer densities in the mouse increase in abundance right after death (0.5 h) and continued to increase, reaching a maximum abundance at 12 h, and then slowly decline (**Fig *11*A**). In the zebrafish, the *Pimr* gene transcripts increased slightly after death (relative to live controls) and abruptly increased after 12 h to maximize at 24 h (**Fig *11*B**). One-way to interpret these phenomena are that the transcripts are depleting the miRNA and/or RBP pools. In response to the decrease, a select group of genes involved in survival and stress compensation were passively transcribed, which accounts for the increases in transcript abundances in our original study.

**Fig 11.**
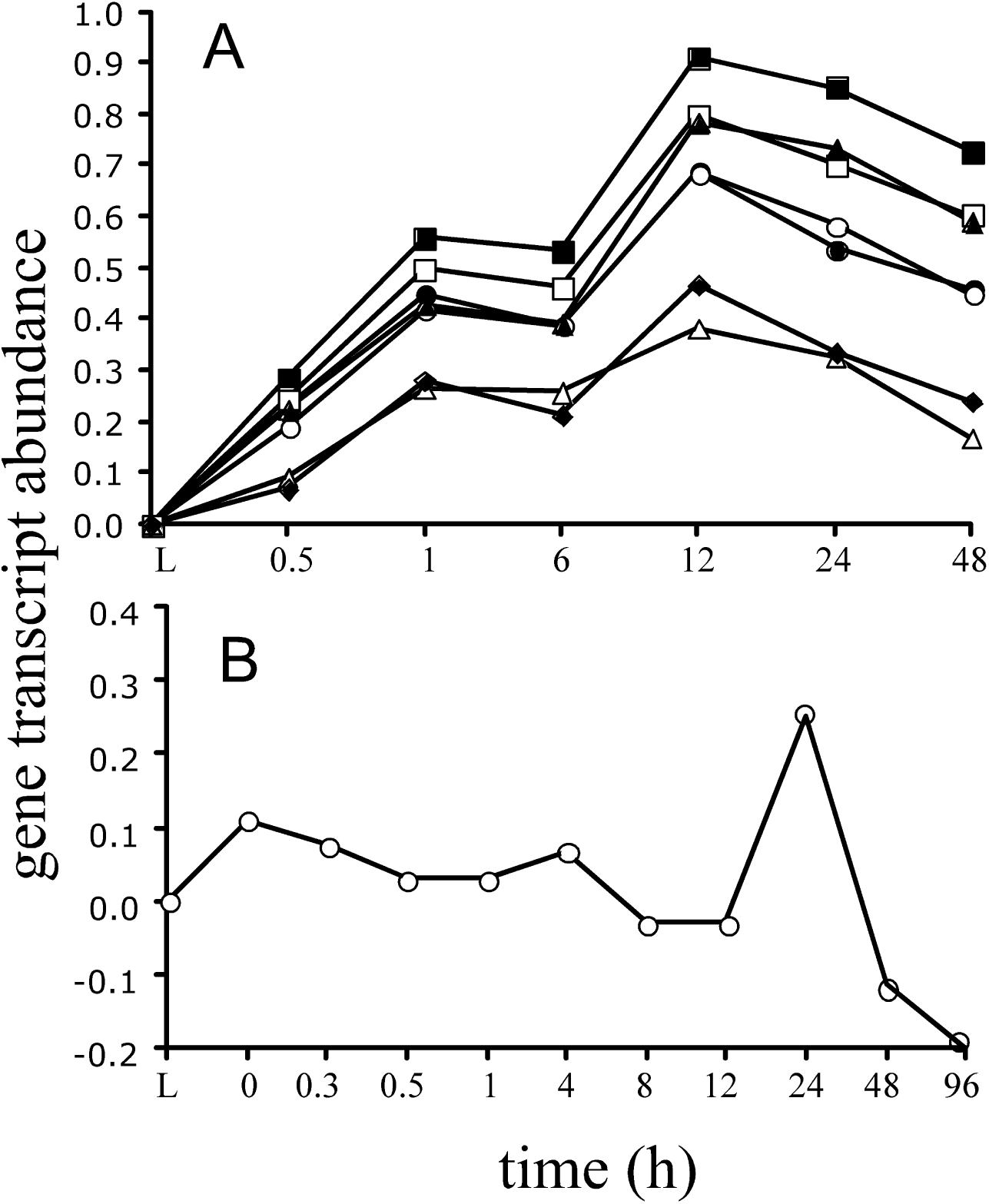
Gene transcript abundances measured by a calibrated microarray [41, 42] (log transformed) by postmortem time. Abundances were normalized to flash frozen live controls (L). Black line, average. (A) Mouse: Open circle, represents Gm11007, Gm2007, Gm4631, Gm14434, Gm2026, Gm14305, Gm14399, Gm14325, Zfp969, Gm4724, Gm14326 transcripts; closed circle, Zfp967, Zfp969, Zfp968; open square, Gm14410; closed square, Gm14305; open triangle, Gm14322; closed triangle, Gm14308; closed diamond, Gm14412. All points are the average of 3 replicates per sample time except the 48 h, which is the average of 2 replicates. (B) Zebrafish: *Pimr* transcript. Each point in the zebrafish represents the average of two individuals per sample time. Data are from ref. [4].

Further support for this scenario comes from the fact that most of the functional genes involved in survival and stress compensation were found in two clusters in the mouse: Groups A and G (59% of the OP) with low mer densities of 0.11 ± 0.12 mers/nt and 0.11 ± 0.05 mers/nt, respectively (**Fig *6***). In the zebrafish, most of the known functional gene transcripts are dispersed into groups A, C, F, K L, M, and N (93% of the OP) (**Fig *5***), which have low mer densities (e.g., 0.10 ± 0.02 mers/nt). It is these genes that might have been passively upregulated due to lack of miRNA and RBPs to prevent them. This scenario makes sense for an evolutionary perspective because post-transcriptional regulation facilitates fast changes in response to stress so that cells can adapt to environmental change.

### Alternative splicing sites might differ under stress

We assumed that the mRNA transcripts downloaded from NCBI represent dominant isoforms one would expect to find in nature. However, a recent study [3] suggests that stress increases the production of different isoforms through alternative splicing. In other words, the composition of the transcripts might change in stressful conditions (i.e., different isoforms are produced). Our analysis did not account for this, however repeating our experiment using next-generation-sequencing methods might indeed provide additional insight into post-transcriptional regulation in postmortem gene expression, which is the focus of our future research.

### Competing endogenous RNA hypothesis

According to the ‘competing endogenous RNA’ hypothesis, all types of RNA transcripts communicate through regulatory-binding sites and it is these interactions that regulate gene expression [39]. The binding of miRNAs to sites represses translation and destabilizes the mRNA, thus having an overall negative regulatory role on gene expression. However, in the case when there is a limited pool of miRNAs to bind the sites or an overabundance of binding sites in transcripts, there is competition between targets to sequester miRNA. Thus, a surplus of binding sites dilutes the miRNA pool and gene expression resumes passively. Pseudogenes (i.e., those resembling known genes but are nonfunctional) as well as other transcripts can dilute the miRNA pool and thereby regulate their availability, and thus have an overall positive regulatory role on gene expression.

Missing from the competing endogenous RNA hypothesis is the role of RBPs to compete with miRNA for regulatory binding sites. The presumed reason for this omission was at the time (i.e., 2011) there was a paucity of information supporting the idea that molecular sponges interact with proteins. However, proof exists today [22]. A recent study reanalyzed high-throughput cross-linking and immunoprecipitation experiments in Human Embryonic Kidney Cells 293 to show that RBPs and miRNA often bind to the same or overlapping regulatory binding sites. The significance of this finding is twofold: (i) it suggests competition among the regulators (RBPs, miRNA, binding sites in different targets) and (ii) it suggests the relative concentrations of the RBPs and miRNAs to the regulatory binding sites might determine a transcript’s fate [40].

A third significant finding from the same study was the introduction of ‘hotspot’ binding sites that have high sequence conservation, accessibility, and enrichment in AU-rich elements (AREs) (i.e., devoid of guanines) and function by favoring competition among regulators [40]. Apparently, target sites outside of hotspots have increased expression levels compared to targets sites within hotspots. Hence ‘hotspots’ are considered functional regulatory elements that provide an extra layer of regulation of post-transcriptional regulatory networks.

## SUMMARY

This is the first study to investigate over-abundant mers in transcriptomic profiles after organismal death and raises interesting questions relative to post-transcriptional regulation and molecular biology.

## ETHICAL STANDARDS

The experiments comply with the current laws of the USA.

## Acknowledgments

We thank Shivani Soni for reviewing earlier versions of the manuscript.

## SUPPLEMENTARY INFORMATION

1. Online Resource_1. Title: Online Resource_1.docx. Description: Proof that using the ‘Chaos Genome Representation’ method to extract mers from the transcript sequences is more practical (computational efficient) than string-based search algorithms.
2. Online Resource_2. Title: Online Resource_2.xlsx. Description: Two sheets in MS Excel file: (i) zebrafish_probes_PM, and (ii) mouse_probes_PM. PM, perfect match probes. Each sheet has two columns: first column is Agilent Probe ID and second column is DNA sequence.
3. Online Resource_3. Title: Online Resource_3.fna. Description: Two columns in text file of the over-abundant pool (OP) for the mouse. One column is Agilent Probe ID linked to Annotated Gene Name and second column is cDNA sequence. Total of 330 rows.
4. Online Resource_4. Title: Online Resource_4.fna. Description: Two columns in text file of the over-abundant pool (OP) for the zebrafish. One column is Agilent Probe ID linked to Annotated Gene Name and second column is cDNA sequence. Total of 230 rows.
5. Online Resource_5. Title: Online Resource_5.fna. Description: Two columns in text file of the control pool (CP) for the mouse. One column is Agilent Probe ID linked to Annotated Gene Name and second column is cDNA sequence. Total of 32611 rows.
6. Online Resource_6. Title: Online Resource_6.fna. Description: Two columns in text file of the control pool (CP) for the zebrafish. One column is Agilent Probe ID linked to Annotated Gene Name and second column is cDNA sequence. Total of 27433 rows.
7. Online Resource_7. Title: Online Resource_7.xls. Description: Two sheets in MS Excel file: (i) zebrafish, and (ii) mouse. Each sheet has 4 columns: first column is string length (strlen) of OP transcript; second column is blank; third column is string length of the corresponding CP transcript; fourth column is string length of corresponding CP2 transcript. Rows 1 to 231 in the zebrafish sheet contain the strlen of 230 transcripts in both OP and CP1 and CP2; rows 233 and 234 contains average and standard deviations of the columns; row 236 contains the two-tailed t-test results for OP vs CP1 and OP vs CP2. Rows 1 to 334 in the mouse sheet contain the strlen of 333 transcripts in both OP and CP1 and CP2; rows 336 and 337 contains average and standard deviations of the columns; row 339 contains the two-tailed t-test results for OP vs CP1 and OP vs CP2.
8. Online Resource_8. Title: Online Resource_8.xlsx. Description: Nine sheets in MS Excel file. The first sheet provides a detailed Readme that describes the sheets. Basically, first column is abundance of mer in OP, second column is average abundance in CP, third column is standard deviation in CP, and remaining 30 columns are abundances of 30 random draws from CP. Rows differ by mer length.
9. Online Resource_9. Title: Online Resource_9.xlsx. Description: Nine sheets in MS Excel file. The first sheet provides a detailed Readme that describes the sheets. Basically, first column is abundance of mer in OP, second column is average abundance in CP, third column is standard deviation in CP, and remaining 30 columns are abundances of 30 random draws from CP. Rows differ by mer length.
10. Online Resource_10. Title: Online Resource_10.xlsx. Description: 11 sheets in MS Excel file. It is similar to the Online Resource_8 file except the raw data is missing to reduce matrix size. The purpose of the sheets is to calculate over- and under-abundant mers that are 5 X the standard deviation of the CP for each mer. The first sheet provides a detailed Readme that describes the sheets. Rows differ by mer length.
11. Online Resource_11. Title: Online Resource_11.xlsx. Description: 11 sheets in MS Excel file. It is similar to the Online Resource_9 file except the raw data is missing to reduce matrix size. The purpose of the sheets is to calculate over- and under-abundant mers that are 5 X the standard deviation of the CP for each mer. The first sheet provides a detailed Readme that describes the sheets. Rows differ by mer length.
12. Online Resource_12. Title: Online Resource_12.xlsx. Description: Multiple sheets in MS Excel file. The first sheet provides a detailed Readme that describes the sheets. Rows differ by mer length. The matrix file consists of 2245 columns and 230 rows.
13. Online Resource_13. Title: Online Resource_13.xlsx. Description: Multiple sheets in MS Excel file. The first sheet provides a detailed Readme that describes the sheets. Rows differ by mer length. The matrix file consists of 5117 columns and 333 rows.
14. Online Resource_14. Title: Online Resource_14.xls. Description: Two sheets in MS Excel file. The first sheet is the collapsed data of the Zebrafish and the second sheet is the collapsed data of the Mouse. Each sheet shows how the data was log normalized for making the heatmaps. The collapsed data was based on two way cluster groups using Wards linkage methods.
15. Online Resource_15. Title: Online Resource_15.xls. Description: Four sheets in MS Excel file. The first sheet provides a detailed Readme that describes the sheets. The second and third sheets have the number of mer hits by transcript sequence length for the zebrafish and mouse. The fourth sheet has the summarize RegRNA2 output for 10 samples.
16. Online Resource_16. Title: Online Resource_16.xls. Description: Two sheets in MS Excel file. The first sheet is the number of mers by region for the zebrafish and the second sheet is the same for the mouse.

